# The clinical drug candidate ebselen attenuates inflammation and promotes microbiome recovery after antibiotic treatment for *Clostridium difficile* infection

**DOI:** 10.1101/827329

**Authors:** Megan Garland, Andrew J Hryckowian, Martina Tholen, Sebastian Loscher, William W. Van Treuren, Kristina Oresic Bender, Justin L Sonnenburg, Matthew Bogyo

## Abstract

*Clostridium difficile* infection (CDI) is an enteric bacterial disease that is increasing in prevalence worldwide. *C. difficile* capitalizes on gut inflammation and microbiome dysbiosis to establish infection, with symptoms ranging from watery diarrhea to toxic megacolon. We recently reported that the safe in human clinical drug candidate ebselen (NCT03013400, NCT01452607, NCT00762671, NCT02603081) has biochemical, cell-based and *in vivo* efficacy against the bacterial toxins of *C. difficile*. Here, we show that ebselen treatment reduces recurrence rates and decreases colitis in a hamster relapse model of CDI. Furthermore, ebselen treatment does not alter microbiome diversity but promotes its recovery back to that of healthy controls after antibiotic-induced dysbiosis in both healthy and *C. difficile*-infected mice. This increased microbiome recovery upon ebselen treatment correlates with a decrease in host-derived inflammatory markers suggesting that the anti-inflammatory properties of ebselen, combined with its anti-toxin function, help to mitigate the major clinical challenges of CDI, including recurrence, microbial dysbiosis, and colitis.

## Introduction

The enteric diarrheal illness caused by *Clostridium difficile* infection (CDI) is an increasing health threat. CDI directly causes 15,000 deaths per year in the US alone and is an exacerbating comorbidity in a further 14,000 US deaths per year (Lessa et al., 2015). CDI is typically incited by antibiotic use, which induces dysbiosis in the commensal microbial communities of the gastrointestinal (GI) tract, allowing this pathogen to thrive. Risk factors such as antibiotic use, proton pump inhibitors (PPIs) and Westernized diets encourage GI inflammation and alter microbial commensals to reduce community robustness required for colonization resistance to enteric pathogens (Desai et al., 2016; Hryckowian et al., 2017; Hryckowian et al., 2018; Sonnenburg et al., 2016; Trifan et al., 2017).

A variety of antibiotics taken for unrelated conditions have been shown to disrupt the GI microbiome in human patients and animal models of disease, causing a GI state that is permissive to *C. difficile* colonization and outgrowth (Ng et al., 2013; Schubert et al., 2014; Schubert et al., 2015). Antibiotic selection for CDI-permissive states is even more concerning given clinical experiments showing that even one dose of antibiotics can cause long-term shifts in the GI microbiome (Zaura et al., 2015) and because the antibiotics vancomycin, metronidazole, and fidaxomicin are the current standard-of-care treatments for CDI (Cohen et al., 2010; McDonald et al., 2018). These antibiotics only achieve a clinical cure 72 – 81% of time (Johnson et al., 2014) and patients diagnosed with CDI for the first time have approximately a 20% chance of recurrence (Kelly, 2012; Lessa et al., 2015). After the first recurrence, risk of subsequent recurrences can be as high as 50% (Kelly, 2012). One study using a murine infection model found that as little as two days after removal of vancomycin treatment, *C. difficile* colonization could be achieved upon re-challenge with the bacterium (Schubert et al., 2015). Clearly, standard of care antibiotic therapy carries the significant drawback that a healthy microbiome that would naturally provide colonization resistance is never allowed to recover. Shifting the microbiome back to a healthy state has been shown to reset CDI-permissive states to one that protects against pathogenic colonization with fecal microbiota transplant (FMT) (Hryckowian et al., 2017; van Nood et al., 2013). This treatment relies on the donation of fecal samples from healthy donors to repopulate the GI tract of patients with severe CDI or multiply-recurrent CDI (Seekatz et al., 2014). While this approach is highly effective, lack of standardization between healthy donors, or, indeed, a mechanistic understanding of what defines a healthy donor sample, leaves many questions and the potential for unforeseen adverse effects with this approach.

In addition to microbial dysbiosis, GI inflammation is hypothesized to play a complex role in CDI-mediated disease (Hryckowian et al., 2017; Shen, 2012). While GI dysbiosis itself is characterized by inflammation and initially creates a permissive environment for *C. difficile* colonization and outgrowth, cytotoxicity mediated by *C. difficile* exotoxins TcdA and TcdB maintains inflammation in the GI that favors an optimal niche for continued survival (Hryckowian et al., 2017).

One alternative paradigm for CDI treatment that could spare the commensal microbiome, mitigate colon pathology, and reduce recurrence is the use of antivirulence agents that directly target the toxin mediators of disease. This approach has been validated by Merck, which recently gained FDA-approval for the monoclonal antibody bezlotoxumab (Zinplava), which targets TcdB (Wilcox, 2015). However, the high cost of monoclonal antibody production and intravenous route of administration likely reserves its use to a select patient population, such as those with severe or multiply recurrent disease, as evidenced by market data indicating that only 5000 – 8000 units have been prescribed per month in 2019 (despite a yearly CDI burden of almost half a million patients annually in the US alone (Lessa et al., 2015)) (Database, 2019). The recent report repurposing the antihelminthic agent niclosamide for CDI by targeting host processes in toxin uptake reporting minimal disruption to the GI microbiome further highlights the importance of effects on the microbiome in the assessment of novel CDI treatments (Tam et al., 2018).

Recently, we reported a small-molecule compound, ebselen, with biochemical, cellular, and *in vivo* efficacy against TcdA and TcdB (Bender et al., 2015). We found that ebselen irreversibly inactivated the cysteine protease domain through modification of the active-site cysteine, though potential other beneficial effects of this compound may also include inactivation of the glucosyltransferase domain (Beilhartz et al., 2016) and anti-inflammatory effects (Antony and Bayse, 2011; Sies, 1993). In a clinically-relevant mouse model of CDI, ebselen attenuated toxin-induced GI pathology to the level of uninfected controls (Bender et al., 2015).

Here, we show that ebselen reduces recurrence rates and decreases colitis in a hamster model of relapsing CDI. To elucidate mechanisms underlying this effect, we analyzed microbial communities and host-derived markers of inflammation. We show that ebselen treatment does not alter the healthy microbiome. Rather, treatment with ebselen promotes microbiome recovery from dysbiosis induced by standard of care antibiotic treatment in healthy and *C. difficile*-infected mice. Ebselen also attenuates host-derived GI inflammation after antibiotic treatment in *C. difficile*-infected mice. These data argue for the rapid clinical advancement of ebselen as a therapy for CDI.

## Results

### Ebselen reduces inflammation and recurrence in a hamster model of relapsing CDI

To elucidate benefits of a small-molecule antivirulence drug in mitigating major clinical challenges in the treatment of CDI, we sought to test the efficacy of ebselen in the acute phase of infection as well as recurrence in a hamster model of relapsing CDI. Classically thought of as the gold-standard animal model of CDI (Best et al., 2012), pre-treatment with antibiotics induces microbial dysbiosis that permits *C. difficile* colonization (Figure 1A). However, hamsters are exquisitely sensitive to CDI and, unlike humans, uniformly succumb to infection within days of *C. difficile* challenge in the absence of treatment, unlike humans (Best et al., 2012; Lessa et al., 2015).

**Figure 1:**
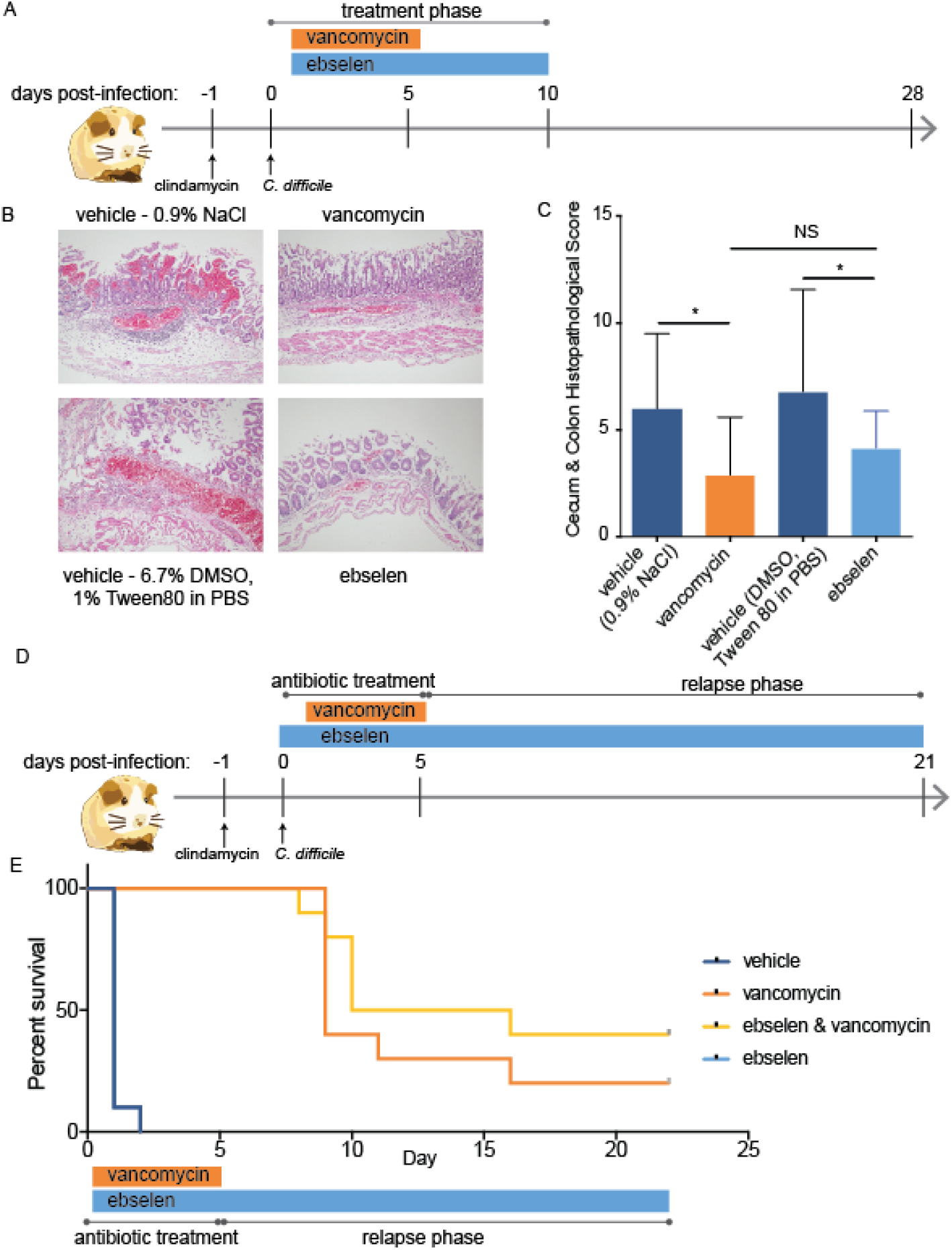
Ebselen reduces inflammation and recurrence in a hamster model of relapsing CDI. A. Schematic of hamster model of CDI. Golden Syrian hamsters were randomly divided into four treatment groups: vehicle 1 (n = 5), vehicle 2 (n = 4), ebselen (n = 10) or vancomycin (n = 10). Dysbiosis was induced with clindamycin on day −1 prior to challenge with BI/NAP1 027 ribotype *C. difficile* (BAA-1805) spores on day 0. Vancomycin was dosed twice daily via oral gavage starting on day 0 for 5 days. Ebselen was dosed twice daily via oral gavage starting on day 0 for 10 days. Hamsters were observed for 28 days post-infection. B. Representative histological images for hamster model of CDI. The cecum and colon were harvested from animals at time of humane sacrifice, death, or on day 28 from surviving animals. Samples were formalin fixed and paraffin embedded for HnE staining and histopathology. C. Histopathology quantification from cecum and colon samples in (B). Slides were scored blinded by a board-certified veterinary pathologist for epithelial damage, vascular congestion and hemorrhage, and inflammation, with statistical analysis by unpaired t-test. * p < 0.05, NS, non-significant. D. Schematic of hamster model of relapsing CDI. Hamsters were randomly divided into four treatment groups of 10 animals each: vehicle, ebselen, vancomycin, or vancomycin + ebselen. Dysbiosis was induced with clindamycin on day −1 prior to challenge with BI/NAP1 027 ribotype *C. difficile* (VA20) spores on day 0. Vancomycin was dosed once daily via oral gavage starting on day 0 for 5 days. Ebselen was dosed twice daily via oral gavage starting on day 0 for the duration of the study. Hamsters were observed for 21 days post-infection. E. Kaplan-Meier curve for relapse hamster model. Hazard ratio of co-treatment with ebselen + vancomycin to vancomycin: 0.537 (95% CI 0.1595 - 1.808) measured by Mantel-Haenszel test.

Using this model, we first asked whether ebselen reduces colitis in the acute phase of infection. Golden Syrian hamsters were pre-treated with clindamycin on day −1 to induce GI microbial dysbiosis, then challenged with a vancomycin-sensitive NAP1/027 strain of *C. difficile*expressing TcdB and TcdA (BAA-1805). Hamsters were orally treated with vancomycin, ebselen, or vehicle and monitored for survival. Infected hamsters treated with ebselen had similar mortality rates as vehicle-treated controls (Supplemental Figure 1A) and did not reduce colony forming units (CFUs) of *C. difficile* spores measured in fecal pellets (Supplemental Figure 1B), consistent with our previous study (Bender et al., 2015). Therefore, ebselen’s lack of effect on reducing mortality rate is likely due to the fact that CDI with toxigenic *C. difficile* is uniformly fatal in hamsters without treatment that significantly reduces CFUs. Regardless, histological analysis of cecum and colon tissue harvested at the time of sacrifice showed a statistically significant decrease in colitis in the ebselen-treated group as compared to vehicle-treated controls as measured by epithelial damage, congestion and hemorrhage and inflammatory cell infiltrate (Figure 1B and 1C). This reduction in the morbidity associated with primary CDI agreed with our results from the murine model (Bender et al., 2015).

Treatment with vancomycin resulted in survival through the treatment period, akin to clinical cure in human patients, but the majority of hamsters eventually experienced a relapse of the infection, leading to death beginning at day 11 post-infection (Supplemental Figure 1A). Though recurrence in human infection does not typically lead to mortality, high recurrence rates are typical of human infection (Lessa et al., 2015). We next asked whether ebselen treatment would result in a reduction of CDI recurrence rates. Using a 027 ribotype strain of *C. difficile* clinically isolated from patients (VA20), we compared *C. difficile* infected hamsters treated with five days of vancomycin with or without concurrent treatment of ebselen, as well as to vehicle- and ebselen-treated control groups (Figure 1D). All hamsters treated with vancomycin survived through the initial infection period (Figure 1E). Hamsters treated with vancomycin began to experience recurrence as measured by mortality around day 9. In this recurrence phase, we observed a decrease in recurrence rates in the group treated with ebselen, with 4 of 10 hamsters surviving to day 21 whereas 2 of 10 survived to day 21 in the vancomycin only group (Figure 1E). This reduction of recurrence rate corresponded to an improved hazard ratio of 0.537 (95% CI 0.1595 - 1.808) in the ebselen and vancomycin treatment group as compared to vancomycin treatment alone. Together, these data show that ebselen treatment alone reduces morbidity in primary CDI as well as decreased recurrence rates in hamsters using clinically-relevant NAP1/027 strains of *C. difficile*, which are associated with increased rates of ileus, toxic megacolon and pseudomembranous colitis and mortality (See et al., 2014).

### Ebselen does not alter the GI microbiome

To dissect mechanisms by which ebselen reduced recurrence rates, we next sought to study major risk factors of CDI and recurrent CDI. Microbial dysbiosis, primarily caused by antibiotic use, appears to be the primary risk factor in CDI (Bignardi, 1998; McDonald et al., 2018). We hypothesized that treatment with our small-molecule anti-virulence agent would not alter the microbiome, while vancomycin would induce dysbiosis characterized by a loss of population diversity.

To profile changes in the GI microbiome during treatment, healthy Swiss Webster mice were orally treated with ebselen, vancomycin, ebselen and vancomycin, or vehicle control for five days (Figure 2A). Fecal samples were collected daily and used for 16S rRNA amplicon analysis. Inter-sample diversity was measured via weighted UniFrac distance between samples and visualized on principle component analysis (PCoA) plots. Two principle components explain 30% of the variance in weighted UniFrac distances between samples (Supplemental Figure 2). As expected, vancomycin treatment was a strong modifier of the mouse microbiome as compared to pre-treatment controls and vehicle-treated mice, with mice in vancomycin or the combination of ebselen and vancomycin treatment groups forming a dysbiotic cluster (Figure 2B, Day 5). Weighted UniFrac distances measured between samples were quantified on day 5 (Figure 2C). Intra-sample variance within each treatment group was small, as indicated by tight clustering of samples on the PCoA plot and minimal UniFrac distance between samples (Figures 2B and 2C). Comparisons between GI microbiomes in the vehicle and ebselen treatment groups showed similar tight clustering, indicating that ebselen treatment did not affect the GI microbiome (Figure 2C). In contrast, comparisons of weighted UniFrac distances between vehicle- and ebselen-treated mice versus the weighted UniFrac distances between vehicle- and vancomycin-treated mice showed a statistically significant difference (Figure 2C, p < 0.0001), highlighting the dramatic difference in effect on the GI microbiome of these two treatments.

**Figure 2:**
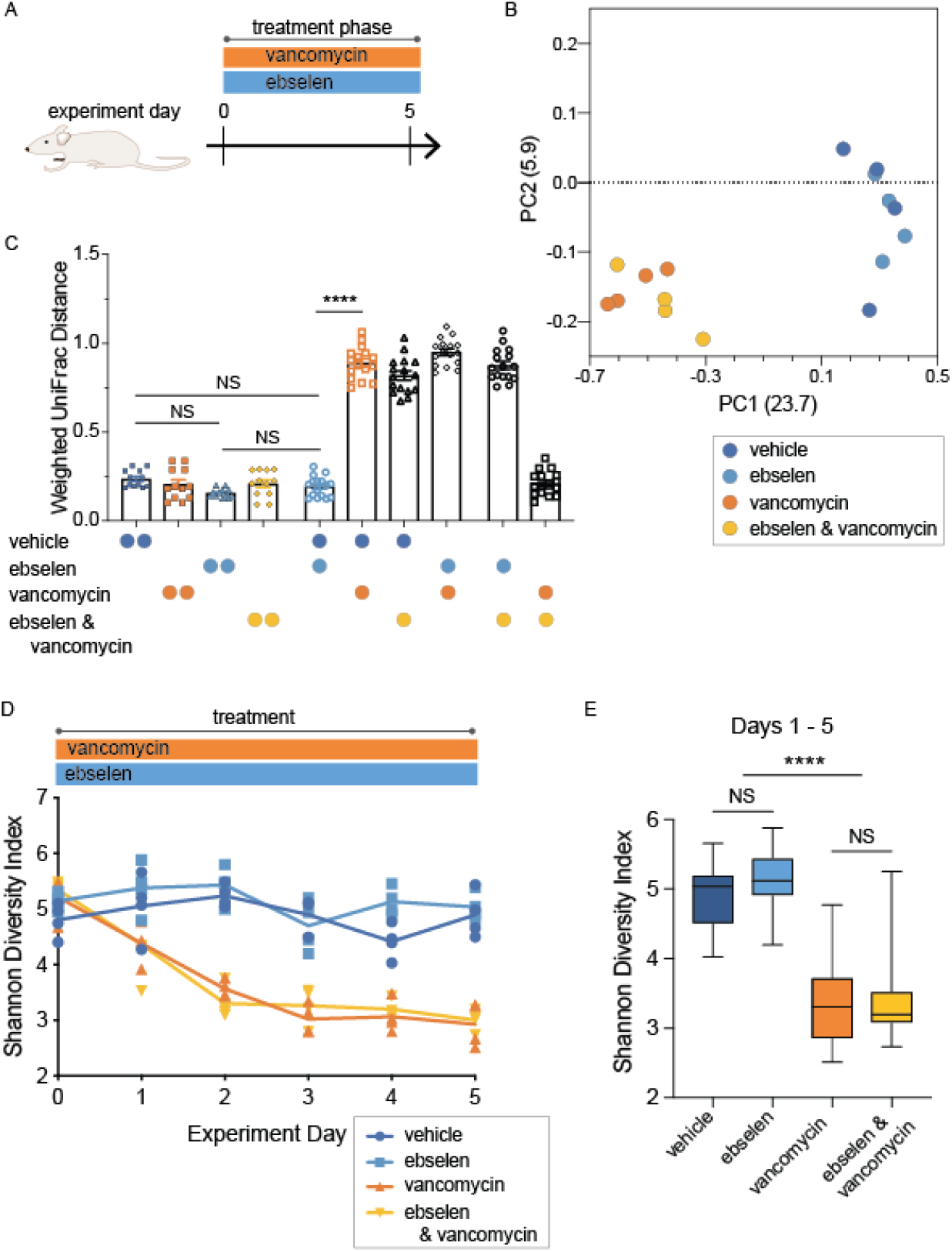
Ebselen does not alter the GI microbiome. A. Conventional Swiss Webster mice were randomly divided into four treatment groups of four animals each treated with one of four treatments: vehicle (6.7% DMSO, 1% Tween80 in PBS), ebselen (100 mg/kg), vancomycin (100 mg/kg), or vancomycin + ebselen (100 mg/kg and 100 mg/kg, respectively). Animals were dosed once daily via oral gavage for five treatments starting on day 0 after sampling. Fecal samples were collected daily for the 5-day experiment. B. Daily fecal samples were analyzed for GI microbial diversity via 16S rRNA amplicon analysis. Principle component analysis (PCoA) plot of weighted UniFrac distances between microbiota of individual mice on day 5. C. Weighted UniFrac distances between microbiota as shown in (B). Quantification represents intra-group measurements (side-by-side spheres, all intra-group comparisons NS) and inter-group comparisons, as indicated by spheres below. Statistics measured via one-way ANOVA with multiple comparisons; **** p < 0.0001. D. Alpha diversity of GI microbial amplicon sequence variants (ASVs) measured over the treatment period via the Shannon diversity index. E. Statistical analysis of microbial diversity shown in (D). Samples representing the treatment period (days 1 – 5) were collapsed for each group. Significance was determined by the Kruskal-Wallis test with the Dunn’s multiple comparison test. NS, non-significant; **** p < 0.0001. ***(148, 964)***

Next, we measured intra-sample Shannon diversity (alpha diversity), showing the richness and abundance of amplicon sequence variants (ASVs) per group per day. Treatment with vancomycin reduced diversity of the gut microbiota when compared to vehicle-treated control (Figure 2D), in accordance with dysbiosis seen in previous studies after vancomycin treatment (Schubert et al., 2015). In contrast, ebselen treatment did not alter the diversity of the microbiome over the five-day treatment period, as compared with vehicle-treated control (Figure 2D). Co-treatment with ebselen and vancomycin was dominated by the vancomycin-induced reduction in unique ASVs over this five-day period. Statistical analysis of microbial diversity collapsed over the five-day treatment period showed no difference in microbial diversity in ebselen-treated versus vehicle-treated controls (Figure 2E). Treatment with vancomycin or vancomycin and ebselen resulted in a statistically significant reduction in diversity (Figure 2E, p < 0.0001). Together, these data show that ebselen does not alter the GI microbiome, while vancomycin treatment induces dysbiosis.

### Ebselen promotes microbiome recovery after antibiotic treatment

We next asked whether treatment with ebselen altered the composition of microbial species during recovery from vancomycin-induced dysbiosis. Healthy Swiss Webster mice were orally treated with ebselen, vancomycin, ebselen and vancomycin co-treatment, or vehicle control for five days (Figure 3A). Fecal samples were collected during the treatment period and subsequent recovery period for a total of 28 days, then analyzed with weighted PCoA analysis of UniFrac distance. Two principle components explain 30% of the variance in weighted UniFrac distances between samples (Supplemental Figure 3). In the treatment phase, microbial communities of the mice treated with vancomycin alone or in combination with ebselen were remodeled, as in Figure 2, to engender a microbiome community composition distinct from vehicle- and ebselen-treated mice (Supplemental Figure 2A), with a similar decrease in species diversity measured by Shannon diversity (Figure 3B).

**Figure 3:**
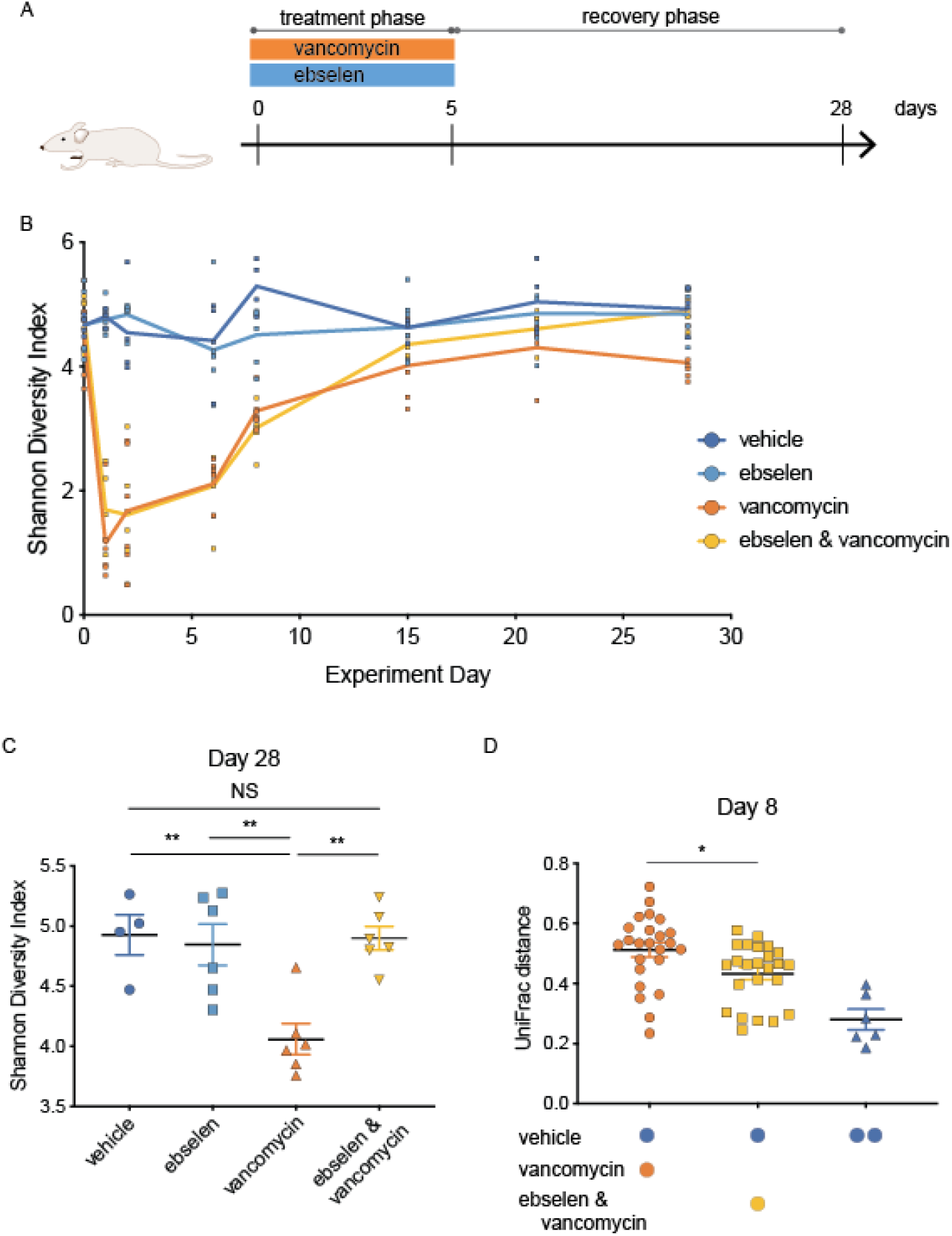
Ebselen promotes microbiome recovery after antibiotic treatment. A. Schematic of experiment. Conventional Swiss Webster mice were randomly divided into four treatment groups of six animals, each treated with one of four treatments: vehicle (6.7% DMSO, 1% Tween80 in PBS), ebselen (100 mg/kg), vancomycin (100 mg/kg), or vancomycin + ebselen (100 mg/kg and 100 mg/kg, respectively). Treatments were dosed once daily via oral gavage for five treatments (treatment phase) starting on day 0 after sampling. B. Fecal samples were analyzed for GI microbial diversity via 16S rRNA amplicon analysis. Alpha diversity of GI microbial amplicon sequence variants (ASVs) were measured over the treatment and recovery phases via Shannon diversity index. C. Statistical analysis of microbial diversity shown in (B) at day 28. Statistical analysis by one-way ANOVA. ** p < 0.01 where indicated, all other comparisons nonsignificant (NS). D. Weighted UniFrac distances between treatment groups measured at day 8. Colored spheres indicate groups from which distances are measured. Statistical analysis via one-way ANOVA, * p < 0.05.

During the recovery phase, Shannon diversity in vancomycin-treated groups increased over time following the vancomycin-induced reduction in diversity during the treatment phase (Figure 3B). However, unique ASVs in vancomycin-treated mice remained significantly statistically lower at day 28 as compared to vehicle-treated controls (Figure 3C, p = 0.0040). These data agree with previous studies highlighting the sustained effect of antibiotic treatment on the microbiome after treatment ends (Dethlefsen and Relman, 2011; Theriot et al., 2014). Intriguingly, however, mice treated with ebselen and vancomycin displayed a full recovery of ASV diversity at day 28 compared to vehicle-treated controls (NS, p = 0.9991), with a statistically significant increase in diversity as compared to vancomycin treatment alone (Figure 3C, p = 0.0020). To compare the rate of recovery, we compared weighted UniFrac distances between vancomycin- and vehicle-treated mice versus ebselen and vancomycin- and vehicle-treated controls during the recovery period. We observed a faster recovery of the antibiotic-induced microbial remodeling in the co-treatment group (Figure 3D). Together, these data show that ebselen treatment mitigates antibiotic-induced microbial dysbiosis by improving diversity and recovery rates back to that of healthy controls.

### Ebselen promotes microbiome recovery and attenuates host-derived GI inflammation in CDI

We next asked whether the ebselen also promoted microbiome recovery after vancomycin use in the context of CDI. Using a clinically-relevant non-fatal mouse model of CDI (Chen et al., 2008), we compared oral treatment with vehicle, ebselen, vancomycin, and ebselen and vancomycin treatment for five days (Figure 4A). Dysbiosis was induced with an antibiotic cocktail in the drinking water followed by orally administered clindamycin, followed by challenge with 10^8^ *C. difficile* vegetative cells (day 0). Animals were treated orally with vehicle, ebselen, vancomycin, and ebselen and vancomycin treatment for five days. Colony-forming units (CFUs) shed in fecal samples on day 1 were enumerated to ensure infection. CFUs were similar in the vehicle- and ebselen-treated animals and undetectable in the vancomycin- and vancomycin and ebselen-treated groups, consistent with the known mechanisms of action of these treatments (Supplemental Figure 4A). Animals were observed for a total of 28 days post *C. difficile* challenge to characterize changes in microbial populations in the pre-infection, acute and recovery phases of infection.

**Figure 4:**
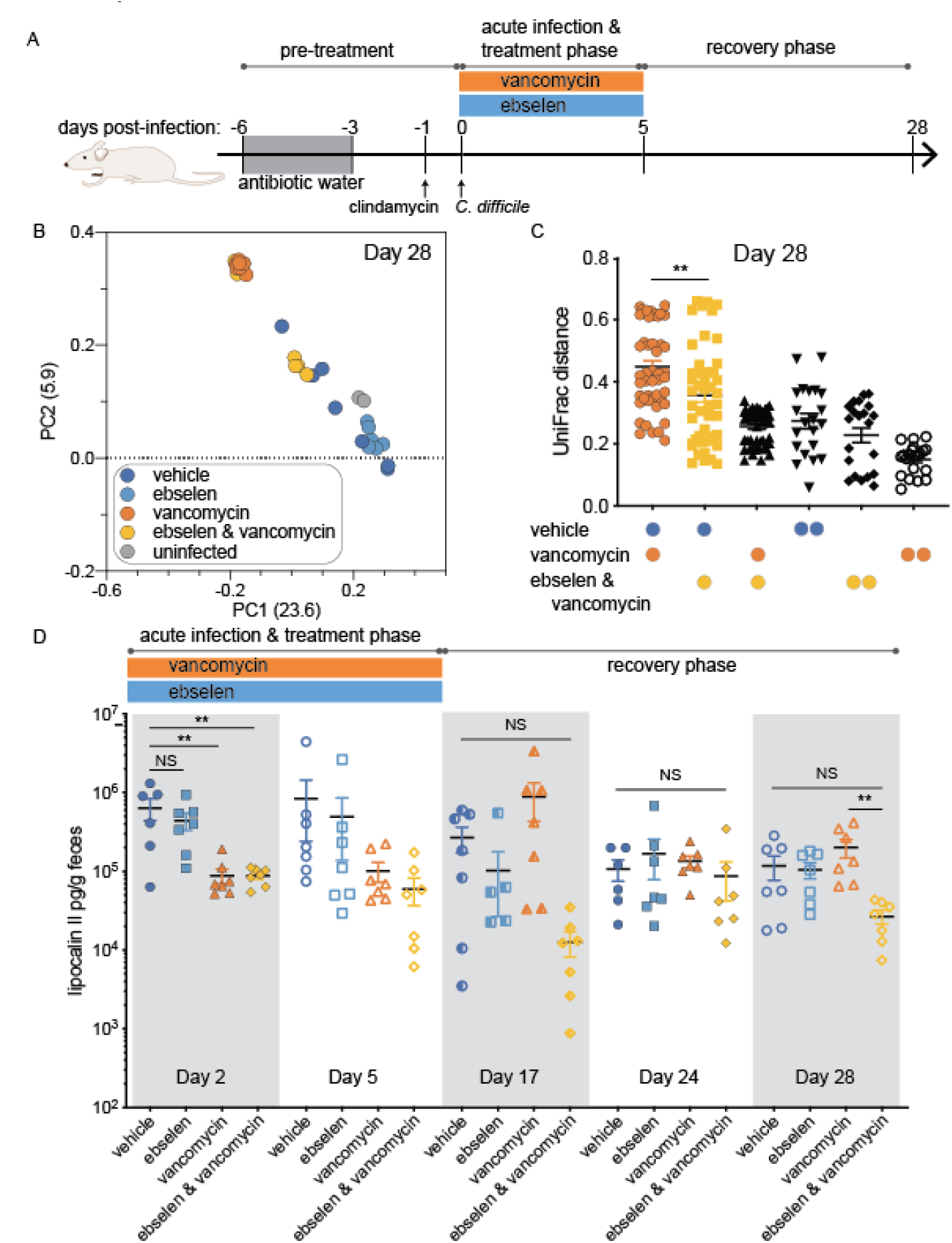
Ebselen attenuates host-derived inflammation and promotes microbiome recovery in a mouse model of CDI. A. Schematic of mouse model of CDI. Dysbiosis was induced in conventional Swiss Webster mice with an antibiotic cocktail in drinking water (kanamycin 0.4 mg/mL, gentamicin 0.035 mg/mL, colistin 850 U/mL, metronidazole (0.215 mg/mL) and vancomycin (0.045 mg/mL)) for three days, beginning six days before inoculation (grey box, pretreatment). Mice were switched to regular water for two days, and then treated with 1 mg of clindamycin via oral gavage on day −1. On day 0, mice were randomly divided into an uninfected control group (n = 5) or four treatment groups of seven animals each treated with one of four treatments: vehicle (6.7% DMSO, 1% Tween80 in PBS), ebselen (100 mg/kg), vancomycin (100 mg/kg), or vancomycin + ebselen (100 mg/kg and 100 mg/kg, respectively). Mice were orally challenged with approximately 10^8^ CFU of *C. difficile* strain 630 (treatment groups) or PBS control. Treatments were dosed once daily via oral gavage for five treatments (acute infection & treatment phase) starting on day 0 after sampling. Fecal samples were collected for 28 days post-infection. B. Fecal samples were analyzed for GI microbial diversity via 16S rRNA amplicon analysis. PCoA plot of weighted UniFrac distances of GI microbiota at day 28. C. Weighted UniFrac distances shown in (B). Colored spheres indicate groups from which distances are measured. Analysis of comparison between weighted UniFrac distances made via one-way ANOVA, ** p < 0.01. D. Fecal lipocalin measured via ELISA for four treatment groups, measured during acute infection & treatment phase (days 2 and 5) and recovery phase (days 17, 24 and 28). Alternating grey and white rectangles delineate different days. Statistical analysis between treatment groups at each day via one-way ANOVA with multiple comparisons. NS, nonsignificant, ** p < 0.01.

We first analyzed the treatment period prior to *C. difficile* challenge (pre-infection) as well as the treatment period during acute infection. Two principle components explain 30% of the variance in weighted UniFrac distances between samples (Supplemental Figure 4B, days −7 to 28). As expected, in the pre-infection phase, pre-treatment with an antibiotic cocktail in the drinking water in addition to orally administered clindamycin led to dysbiosis of microbial communities (Supplemental Figure 4B, Day −7 and day 0) and a decrease in diversity (Supplemental Figure 4C, acute infection and treatment phase). In the acute infection and treatment phase (day 0 – 5), the composition of the GI microbiome appeared to be dominated by treatment rather than *C. difficile* challenge, as treatment groups diverged during this period (Supplemental Figure 4B). GI microbial communities of mice in groups treated with vehicle or ebselen recovered, returning to the cluster containing pre-treatment controls (Supplemental Figure 4B, Day 0 and Day 5). In contrast, the microbial communities of mice treated with vancomycin or ebselen and vancomycin remained clustered at two dysbiotic states (Supplemental Figure 4B, Day 0 and Day 5).

We next analyzed differences between groups in the recovery phase of infection after discontinuation of treatment. At day 28, GI microbial communities in 4 of the 7 mice treated with ebselen and vancomycin recovered, while all of the vancomycin-treated mice remained dysbiotic (Figure 4B). UniFrac distances between two groups, vancomycin-to vehicle-treated mice or ebselen and vancomycin-to vehicle-treated groups at day 28, showed that treatment with ebselen resulted in a more complete recovery towards vehicle-treated controls (Figure 4C). These data suggest that ebselen treatment can compensate for vancomycin-induced dysbiosis by promoting recovery of the microbiome after antibiotic treatment.

Given the interplay between host-derived inflammation and pro-inflammatory microbes, we next sought to measure levels of inflammation in the GI during the acute phase of infection as well as in the recovery phase out to day 28. We hypothesized that ebselen- and cotreated mice would display lower levels of inflammation. We measured levels of fecal lipocalin-2 (NGAL), a neutrophil-derived marker associated with inflammation that has been shown to be elevated in enteric pathogen-susceptible mice as well as in animal models of irritable bowel disease (Chassaing et al., 2012; Desai et al., 2016; Studer et al., 2016). In the acute phase (day 2) we observed a statistically significant decrease in lipocalin-2 with vancomycin- and cotreatment as compared to vehicle-treated controls, consistent with suppression of CDI (Figure 4D). While ebselen alone was shown to reduce inflammatory markers in histological analysis (Bender et al., 2015), the reduction in lipocalin-2 was not statistically significant (Figure 4D, day 2 post-infection). Interestingly, in the recovery phase, vancomycin-treated mice experienced a spike in lipocalin-2 levels similar to those seen in the initial infection with vehicle treatment, which may reflect an ongoing inflammatory response in response to continued gut dysbiosis (Figure 4D, day 17 post-infection). In contrast, host levels of inflammation with ebselen and vancomycin co-treatment did not experience post-infection inflammation and achieved a statistically significant decrease in lipocalin-2 levels compared to vancomycin treatment alone at day 28 (Figure 4D, p < 0.01). These data show that, although vancomycin is initially protective against large increases of host-derived inflammation due to primary CDI, treatment creates increases in GI inflammation in the post-infection period, which may lead to states permissive to CDI recurrence as seen in previous reports (Warren et al., 2013). However, the addition of ebselen to vancomycin treatment yields the benefits of vancomycin by reducing host inflammation in primary CDI, but also protects against ongoing inflammation in the postinfection period.

## Discussion

Here we report that ebselen, a small-molecule anti-virulence agent with a clean safety profile in human clinical trials, reduces inflammation and recurrence in a hamster model of CDI. Our results in murine models of antibiotic-induced dysbiosis and CDI suggest that ebselen’s effect on the microbiome is mediated by promoting recovery after vancomycin-induced dysbiosis and dampening GI inflammation.

Our results indicating that ebselen improves microbiome recovery after standard of care antibiotic treatment is intriguing because to our knowledge, all other reported therapies that improve microbial diversity are microbiota-based therapies such as SER-109 (Khanna et al., 2016) or FMT. This suggests that application of our small-molecule compound may also reduce recurrence rates in humans similar to the results achieved in the hamster model of CDI without the risks associated with treating complex mixtures of microbes (Administration, 2019; Tran et al., 2018).

Recent studies analyzing perturbations to the microbiome with antibiotic use have suggested a stability landscape framework (Shaw et al., 2019) in which antibiotics can serve as a perturbing stimulus that causes the GI microbiome to shift to an alternate equilibrium. In our analysis of the mouse microbiome in the context of CDI, we observed two distinct dysbiotic states in our PCoA plot (see Supplemental Figure 4B). Results at day 28 showed that all vancomycin-treated mice had a similar microbiome while the GI microbiomes of the majority of the mice co-treated with vancomycin and ebselen more closely resembled controls. Application of the stability framework yields the intriguing possibility that vancomycin treatment leads primarily to an alternative stable but dysbiotic state, and that co-treatment with ebselen induces a shift of the microbiome from this stable state back to that of healthy controls.

Given the intriguing differences in alpha and beta diversity we observed between vancomycin treatment with and without ebselen, we searched for specific ASVs with increased or decreased abundance that may account for the changes observed. While species were identified in some datasets, we were unable to identify statistically significant differences that were consistent across all experiments. This is likely due to multiple reasons. First, changes in single species may be less important than the global functional output of the microbiome as a whole, for example, creating metabolic outputs that are pro-versus anti-inflammatory. Second, many studies have identified large differences in the ‘healthy’ microbiome (Lloyd-Price et al., 2016), creating a challenge for generalizing specific compositions of species across studies.

Several caveats exist in the interpretation of this study. First, the hamster relapse model failed to achieve a statistically significant decrease in recurrence rates with ebselen treatment although the hazard ratio and median survival was improved relative to vancomycin alone (13 days with ebselen and vancomycin versus 9 days with vancomycin alone). This is likely due in part to technical limitations in increasing the number of animals per group (n = 10) while comparing multiple treatments. Additionally, the Kaplan-Meier curves cross early in relapse (at day 9), which mathematically increases the p value.

Extensive work has shown that ebselen has antioxidant properties, raising the possibility that some beneficial dampening of the inflammatory response in these models was due to a generic decrease in GI oxidative stress rather than a neutralization of toxin-induced inflammation. Regardless of the exact mechanism of inflammatory reduction seen in these animals, it remains a positive effect. Further, we measured secondary measures of inflammation, i.e. secreted proteins from inflammatory infiltrates in response to bacterial infection, indicating that, regardless of the initial source of decreased inflammation, positive feedback from the host and microbiome sides led to an overall decrease of inflammation in the GI tract. Ultimately, mechanistic understanding of the generic antioxidant effect of ebselen to its ability to reduce colonic inflammation would require development of an equally potent antitoxin, but non-antioxidant small-molecule compound to compare in animal models head to head.

In summary, infection with *C. difficile* involves a complex interplay between the host, the GI microbiome, and external perturbances to yield a state permissive to survival for this opportunistic enteric pathogen. Together with our previous work, we show that treatment with the orally administered compound ebselen reduces short-term colitis caused by CDI as well as long-term sequelae of recurrent disease and a persistent dysbiotic state due to repeated courses of antibiotics. This work suggests three primary treatment options for advancement into the clinic: as a monotherapy for mild disease, in combination with antibiotics for more severe infections to reduce reoccurrence, and finally, as a primary preventative treatment in combination with antibiotic therapy for patients at high risk of CDI.

## Materials and Methods

### Bacterial strains and culture conditions

Murine experiments: *C. difficile* strain 630 was used in all murine animal experiments. Frozen stocks were cultured on CDMN agar plates (*C. difficile* agar base (Oxoid CM0601) supplemented with 7% (v/v) defibrinated horse blood (Lampire Biological Laboratories), 32 mg/L moxalactam (Santa Cruz Biotechnology), and 12 mg/L norfloxacin (Sigma-Aldrich) in an anaerobic chamber (Coy) at 37 °C for 24 hours. Single colonies were picked and grown anaerobically for 16 – 18 hours at 37 °C to saturation in reinforced Clostridial medium (RCM, Oxoid) for inoculation of mice. For quantification of *C. difficile* CFUs, 1 μL of feces was serially diluted in PBS and plated onto CDMN plates and incubated for 18 – 24 hours anaerobically before enumeration.

Hamster model of CDI: *C. difficile* BI/NAP1 027 ribotype strain (BAA-1805) was used for the hamster model of CDI. Frozen stocks were cultured anaerobically on a blood agar plate for 5 days at 37 °C. Colonies were transferred to PBS and heated to 70 °C for 30 minutes to kill vegetative cells. The heated culture were pelleted by centrifugation, 3300 x g for 15 minutes, then resuspended in cold PBS. The culture was diluted in PBS to obtain the estimated inoculum sizes of 2.0 x 10^5^ spores/mL. Final colony counts were determined by plating serial dilutions onto CCFA-HT plates and incubated for 5 days incubation before enumeration.

Hamster model of relapsing CDI: *C. difficile* BI/NAP1 027 ribotype strain, binary toxin positive (VA20) was used for the hamster model of relapsing CDI. Frozen stocks were cultured anaerobically and diluted to obtain the estimated inoculum of 10^2^ spores per animal in 0.5 mL.

### Mouse model of healthy mice

Age-and sex-matched conventional Swiss-Webster mice (SWEF, Taconic) ages 7 to 16 weeks, fed standard chow, with an average weight of 30 g were randomly divided into one of four treatment groups: ebselen (100 mg/kg), vancomycin (100 mg/kg), ebselen (100 mg/kg) and vancomycin (100 mg/kg) co-treatment, or vehicle control. Treatments were administered via daily oral gavage in 200 μL final volume in vehicle (6.7% DMSO, 1% Tween-80 in PBS) for 5 days, starting on day 0. Fecal samples were collected daily before oral gavage treatments then every 2-4 days for 14- or 28-days total. Mice were sacrificed according to the guidelines on humane termination after 14- or 28-day experiments.

### Mouse model of CDI

All animal experiments were conducted in accordance with APLAC protocols approved by the Stanford University Institutional Animal Care and Use Committee (IACUC). Age-and sex-matched conventional Swiss-Webster mice (SWEF, Taconic) ages 7 to 16 weeks, fed standard chow, were pretreated with an antibiotic cocktail (kanamycin (0.4 mg/mL), gentamycin (0.035 mg/mL), colistin (850 U/mL), metronidazole (0.215 mg/mL) and vancomycin (0.045 mg/mL)) in drinking water for 3 days, starting 6 days before inoculation as previously reported (Chen 2008). Mice were switched to regular water for 2 days, and then administered 1 mg of clindamycin via oral gavage 1 day before *C. difficile* (strain 630) inoculation (approximately 2.2 × 10^8^ CFU from overnight cultures via oral gavage). Mice with an average weight of 30 g were randomly divided into one of four treatment groups: ebselen (100 mg/kg), vancomycin (100 mg/kg), ebselen (100 mg/kg) and vancomycin (100 mg/kg) co-treatment, or vehicle control. Treatments were administered via daily oral gavage in 200 μL final volume in vehicle (6.7% DMSO, 1% Tween-80 in PBS) for 5 days, starting with the first dose 2 hours prior to *C. difficile* challenge on day 0. Fecal samples were collected during pre-treatment, treatment, and recovery phases for subsequent 16s rRNA amplicon sequencing. Mouse feces on day 1 was additionally used for CFU counts. Mice were sacrificed according to the guidelines on humane termination on day 28 of the experiment. Colon tissues were collected for histological analysis.

### 16S rRNA amplicon sequencing

Total DNA was extracted from frozen fecal pellets using the PowerSoil-htp 96 well DNA isolation Kit (MoBio) or the DNeasy PowerSoil HTP 96 Kit (Qiagen). Barcoded primes were used to amplify the V3-V4 region of the 16S rRNA gene from extracted bacterial DNA using primers 515fB and 806rB via PCR (EMP). PCR clean-up was performed with UltraClean PCR Clean-Up Kit (MoBio) or UltraClean 96 PCR Cleanup Kit (Qiagen) before quantification of amplicon products with Quant-iT dsDNA Assay Kit (Thermo Fisher). Amplicons were pooled by adding 100 ng of each product. Illumina MiSeq paired-end reads runs were performed, with experiments divided among runs such that no more than 384 samples were read per run.

### 16S rRNA amplicon sequencing and ASV picking methods

Each sequencing run was independently demultiplexed using QIIME 1.9.1 script ‘split_libraries_fastq.py’. Reads were eliminated only if the index (barcode) read contained more than 1 error. Each sequencing run was then passed through the default DADA2 v 1.6.0 (Callahan et al., 2016) pipeline. Forward and reverse reads were truncated at ((200, 130), (200, 180), (180,150)) for experiments bhffl, bhf86, and exp3 respectively. After ASV calling, ASV tables were merged with DADA2 command ‘mergeSequenceTables’, and bimera’s eliminated with DADA2 command ‘removeBimeraDenovo’. The remaining ASV sequences were assigned taxonomy using the script ‘assign_taxonomy’ with defaults in QIIME 1.9.1. Features with an ‘unassigne? taxonomy were eliminated and the remaining ASV sequences were aligned using PyNAST and QIIME 1.9.1 script ‘align_seqs.py’. A phylogenetic tree was constructed from the aligned sequences using QIIME 1.9.1 script ‘make_phylogeny.py’ with default settings. We used ASV tables rarefied to 4,000 in this study, facilitating inter-run comparisons.

### Lipocalin II ELISA

ELISA was performed with using the mouse Lipocalin-2/NGAL DuoSet ELISA kit (R&D Biosystems) following manufacturer instructions, with samples prepared as reported previously with few modifications (Desai 2016). Briefly, frozen fecal samples were thawed on wet ice, with 5 – 10 mg of sample weighed into fresh Eppendorf tubes. Samples were resuspended in cold, sterile PBS to a concentration of 1 mg feces to 100 μL PBS. Samples were mechanically disrupted and vigorously vortexed to resuspend samples homogenously. For ELISA measurements, homogenized samples were diluted 1:10 in a mixture of water and kit-supplied Reagent Diluent 2 (R&D Biosystems) to a final concentration of 1X Reagent Diluent 2 and 0.1 mg fecal sample in the 100 μL used for ELISA measurements.

### Hamster model of CDI

This experiment was conducted in accordance with APLAC protocols approved by the Eurofins Panlabs Taiwan and Stanford University Institutional Animal Care and Use Committees (IACUCs). Male Golden Syrian hamsters weighing 90 ± 10 g were randomly divided into four treatment groups: vehicle 1 (n = 5), vehicle 2 (n = 4), ebselen (n = 10) or vancomycin (n = 10).

Dysbiosis was induced with clindamycin (50 mg/kg, subcutaneous) on day −1. Hamsters were orally challenged with 1.98 x 10^5^ spores/animal for LD_90-100_ infection of a *C. difficile* BI/NAP1 027 ribotype strain (BAA-1805) on day 0. Ebselen (100 mg/kg formulated in 6.7% DMSO, 0.1% Tween80 in PBS), vancomycin (10 mg/kg formulated in 0.9% NaCl), vehicle 1 (6.7% DMSO, 0.1% Tween80 in PBS) or vehicle 2 (0.9% NaCl) was administered orally twice per day (BID) for 5 (vancomycin, vehicle 2) or 10 days (ebselen, vehicle 1) starting 16 hours after inoculation. Mortality was observed for 28 days in all groups. Stools were collected on day 3 to measure spore counts in surviving animals. The cecum and colon were harvested from animals at time of humane sacrifice or death, or on day 28 from surviving animals. Samples were formalin fixed and paraffin embedded for HnE staining and histopathology. Slides were scored blinded by a veterinary pathologist for epithelial damage, vascular congestion and hemorrhage, and inflammation as described previously (Kokkotou et al., 2008; Pawlowski et al., 2010).

### Hamster model of relapsing CDI

This experiment was conducted in accordance with APLAC protocols approved by the University of North Texas Health Science Center and Stanford University Institutional Animal Care and Use Committees (IACUCs). Male Golden Syrian hamsters (80-100 g) were randomly divided into four treatment groups of 10 animals each: vehicle (6.7% DMSO, 1% Tween80 in PBS), ebselen (100 mg/kg), vancomycin (10 mg/kg), or vancomycin + ebselen (10 mg/kg and 100 mg/kg, respectively). Dysbiosis was induced with clindamycin (10 mg/kg, subcutaneous) on day −1. Hamsters were orally challenged with 10^2^ spores of a C. difficile BI/NAP1 027 ribotype strain (VA20) on day 0. Treatment with ebselen (100 mg/kg) was administered twice daily via oral gavage for 21 days, beginning on day 0 two hours prior to *C. difficile* challenge. Vancomycin (10 mg/kg) was administered via oral gavage once daily for 5 days, beginning 16 hours after *C. difficile* challenge. Vehicle (6.7% DMSO, 0.1% Tween80 in PBS) was administered twice daily via oral gavage for 5 days. Mortality was observed for 21 days in all groups.

### Statistical methods

Alpha and beta diversity were computed using QIIME 1.9.1 (‘alpha_diversity_through_plots.py’, ‘beta_diversity_through_plots.py’ and ‘core_diversity_analyses.py’). Statistical analyses (Mantel-Haenszel test, t-test, ANOVA, Kruskal-Wallis test with the Dunn’s multiple comparison test, as indicated in figure legends) were performed using the Prism 7 software package (GraphPad software).

## References

Administration, U.F.a.D. (2019). Important Safety Alert Regarding Use of Fecal Microbiota for Transplantation and Risk of Serious Adverse Reactions Due to Transmission of Multi-Drug Resistant Organisms, U. Fda, ed. (fda.gov).

Antony, S., and Bayse, C.A. (2011). Modeling the mechanism of the glutathione peroxidase mimic ebselen. Inorganic chemistry 50, 12075–12084.

Beilhartz, G.L., Tam, J., Zhang, Z., and Melnyk, R.A. (2016). Comment on “A small-molecule antivirulence agent for treating Clostridium difficile infection”. Science translational medicine 8, 370tc372.

Bender, K.O., Garland, M., Ferreyra, J.A., Hryckowian, A.J., Child, M.A., Puri, A.W., Solow-Cordero, D.E., Higginbottom, S.K., Segal, E., Banaei, N., et al. (2015). A small-molecule antivirulence agent for treating Clostridium difficile infection. Science translational medicine 7, 306ra148.

Best, E.L., Freeman, J., and Wilcox, M.H. (2012). Models for the study of Clostridium difficile infection. Gut microbes 3, 145–167.

Bignardi, G.E. (1998). Risk factors for Clostridium difficile infection. J Hosp Infect 40, 1–15.

Callahan, B.J., McMurdie, P.J., Rosen, M.J., Han, A.W., Johnson, A.J., and Holmes, S.P. (2016). DADA2: High-resolution sample inference from Illumina amplicon data. Nat Methods 13, 581–583.

Chassaing, B., Srinivasan, G., Delgado, M.A., Young, A.N., Gewirtz, A.T., and Vijay-Kumar, M. (2012). Fecal lipocalin 2, a sensitive and broadly dynamic non-invasive biomarker for intestinal inflammation. PLoS One 7, e44328.

Chen, X., Katchar, K., Goldsmith, J.D., Nanthakumar, N., Cheknis, A., Gerding, D.N., and Kelly, C.P. (2008). A mouse model of Clostridium difficile-associated disease. Gastroenterology 135, 1984–1992.

Cohen, S.H., Gerding, D.N., Johnson, S., Kelly, C.P., Loo, V.G., McDonald, L.C., Pepin, J., Wilcox, M.H., Society for Healthcare Epidemiology of, A., and Infectious Diseases Society of, A. (2010). Clinical practice guidelines for Clostridium difficile infection in adults: 2010 update by the society for healthcare epidemiology of America (SHEA) and the infectious diseases society of America (IDSA). Infect Control Hosp Epidemiol 31, 431–455.

Database, S.H. (2019). Zinplava - US Estimates.

Desai, M.S., Seekatz, A.M., Koropatkin, N.M., Kamada, N., Hickey, C.A., Wolter, M., Pudlo, N.A., Kitamoto, S., Terrapon, N., Muller, A., et al. (2016). A Dietary Fiber-Deprived Gut Microbiota Degrades the Colonic Mucus Barrier and Enhances Pathogen Susceptibility. Cell 167, 1339–1353 e1321.

Dethlefsen, L., and Relman, D.A. (2011). Incomplete recovery and individualized responses of the human distal gut microbiota to repeated antibiotic perturbation. Proc Natl Acad Sci U S A 108 Suppl 1, 4554–4561.

Hryckowian, A.J., Pruss, K.M., and Sonnenburg, J.L. (2017). The emerging metabolic view of Clostridium difficile pathogenesis. Curr Opin Microbiol 35, 42–47.

Hryckowian, A.J., Van Treuren, W., Smits, S.A., Davis, N.M., Gardner, J.O., Bouley, D.M., and Sonnenburg, J.L. (2018). Microbiota-accessible carbohydrates suppress Clostridium difficile infection in a murine model. Nat Microbiol 3, 662–669.

Johnson, S., Louie, T.J., Gerding, D.N., Cornely, O.A., Chasan-Taber, S., Fitts, D., Gelone, S.P., Broom, C., Davidson, D.M., and Polymer Alternative for, C.D.I.T.i. (2014). Vancomycin, metronidazole, or tolevamer for Clostridium difficile infection: results from two multinational, randomized, controlled trials. Clinical infectious diseases: an official publication of the Infectious Diseases Society of America 59, 345–354.

Kelly, C.P. (2012). Can we identify patients at high risk of recurrent Clostridium difficile infection? Clinical microbiology and infection: the official publication of the European Society of Clinical Microbiology and Infectious Diseases 18 Suppl 6, 21–27.

Khanna, S., Pardi, D.S., Kelly, C.R., Kraft, C.S., Dhere, T., Henn, M.R., Lombardo, M.J., Vulic, M., Ohsumi, T., Winkler, J., et al. (2016). A Novel Microbiome Therapeutic Increases Gut Microbial Diversity and Prevents Recurrent Clostridium difficile Infection. J Infect Dis 214, 173–181.

Kokkotou, E., Moss, A.C., Michos, A., Espinoza, D., Cloud, J.W., Mustafa, N., O’Brien, M., Pothoulakis, C., and Kelly, C.P. (2008). Comparative efficacies of rifaximin and vancomycin for treatment of Clostridium difficile-associated diarrhea and prevention of disease recurrence in hamsters. Antimicrob Agents Chemother 52, 1121–1126.

Lessa, F.C., Mu, Y., Bamberg, W.M., Beldavs, Z.G., Dumyati, G.K., Dunn, J.R., Farley, M.M., Holzbauer, S.M., Meek, J.I., Phipps, E.C., et al. (2015). Burden of Clostridium difficile infection in the United States. The New England journal of medicine 372, 825–834.

Lloyd-Price, J., Abu-Ali, G., and Huttenhower, C. (2016). The healthy human microbiome. Genome Med 8, 51.

McDonald, L.C., Gerding, D.N., Johnson, S., Bakken, J.S., Carroll, K.C., Coffin, S.E., Dubberke, E.R., Garey, K.W., Gould, C.V., Kelly, C., et al. (2018). Clinical Practice Guidelines for Clostridium difficile Infection in Adults and Children: 2017 Update by the Infectious Diseases Society of America (IDSA) and Society for Healthcare Epidemiology of America (SHEA). Clinical infectious diseases: an official publication of the Infectious Diseases Society of America 66, 987–994.

Ng, K.M., Ferreyra, J.A., Higginbottom, S.K., Lynch, J.B., Kashyap, P.C., Gopinath, S., Naidu, N., Choudhury, B., Weimer, B.C., Monack, D.M., et al. (2013). Microbiota-liberated host sugars facilitate post-antibiotic expansion of enteric pathogens. Nature 502, 96–99.

Pawlowski, S.W., Calabrese, G., Kolling, G.L., Platts-Mills, J., Freire, R., AlcantaraWarren, C., Liu, B., Sartor, R.B., and Guerrant, R.L. (2010). Murine model of Clostridium difficile infection with aged gnotobiotic C57BL/6 mice and a BI/NAP1 strain. J Infect Dis 202, 1708–1712.

Schubert, A.M., Rogers, M.A., Ring, C., Mogle, J., Petrosino, J.P., Young, V.B., Aronoff, D.M., and Schloss, P.D. (2014). Microbiome data distinguish patients with Clostridium difficile infection and non-C. difficile-associated diarrhea from healthy controls. MBio 5, e01021–01014.

Schubert, A.M., Sinani, H., and Schloss, P.D. (2015). Antibiotic-Induced Alterations of the Murine Gut Microbiota and Subsequent Effects on Colonization Resistance against Clostridium difficile. MBio 6, e00974.

See, I., Mu, Y., Cohen, J., Beldavs, Z.G., Winston, L.G., Dumyati, G., Holzbauer, S., Dunn, J., Farley, M.M., Lyons, C., et al. (2014). NAP1 strain type predicts outcomes from Clostridium difficile infection. Clinical infectious diseases: an official publication of the Infectious Diseases Society of America 58, 1394–1400.

Seekatz, A.M., Aas, J., Gessert, C.E., Rubin, T.A., Saman, D.M., Bakken, J.S., and Young, V.B. (2014). Recovery of the gut microbiome following fecal microbiota transplantation. MBio 5, e00893–00814.

Shaw, L.P., Bassam, H., Barnes, C.P., Walker, A.S., Klein, N., and Balloux, F. (2019). Modelling microbiome recovery after antibiotics using a stability landscape framework. ISME J 13, 1845–1856.

Shen, A. (2012). Clostridium difficile toxins: mediators of inflammation. Journal of innate immunity 4, 149–158.

Sies, H. (1993). Ebselen, a selenoorganic compound as glutathione peroxidase mimic. Free radical biology & medicine 14, 313–323.

Sonnenburg, E.D., Smits, S.A., Tikhonov, M., Higginbottom, S.K., Wingreen, N.S., and Sonnenburg, J.L. (2016). Diet-induced extinctions in the gut microbiota compound over generations. Nature 529, 212–215.

Studer, N., Desharnais, L., Beutler, M., Brugiroux, S., Terrazos, M.A., Menin, L., Schurch, C.M., McCoy, K.D., Kuehne, S.A., Minton, N.P., et al. (2016). Functional Intestinal Bile Acid 7alpha-Dehydroxylation by Clostridium scindens Associated with Protection from Clostridium difficile Infection in a Gnotobiotic Mouse Model. Front Cell Infect Microbiol 6, 191.

Tam, J., Hamza, T., Ma, B., Chen, K., Beilhartz, G.L., Ravel, J., Feng, H., and Melnyk, R.A. (2018). Host-targeted niclosamide inhibits C. difficile virulence and prevents disease in mice without disrupting the gut microbiota. Nat Commun 9, 5233.

Theriot, C.M., Koenigsknecht, M.J., Carlson, P.E., Jr., Hatton, G.E., Nelson, A.M., Li, B., Huffnagle, G.B., J, Z.L., and Young, V.B. (2014). Antibiotic-induced shifts in the mouse gut microbiome and metabolome increase susceptibility to Clostridium difficile infection. Nat Commun 5, 3114.

Tran, V., Phan, J., Nulsen, B., Huang, L, Kaneshiro, M., Weiss, G., Ho, W., Sack, J., Ha, C., Uslan, D., et al. (2018). Severe Ileocolonic Crohn’s Disease Flare Associated with Fecal Microbiota Transplantation Requiring Diverting Ileostomy. ACG Case Rep J 5, e97.

Trifan, A., Stanciu, C., Girleanu, I., Stoica, O.C., Singeap, A.M., Maxim, R., Chiriac, S.A., Ciobica, A., and Boiculese, L. (2017). Proton pump inhibitors therapy and risk of Clostridium difficile infection: Systematic review and meta-analysis. World J Gastroenterol 23, 6500–6515.

van Nood, E., Vrieze, A., Nieuwdorp, M., Fuentes, S., Zoetendal, E.G., de Vos, W.M., Visser, C.E., Kuijper, E.J., Bartelsman, J.F., Tijssen, J.G., et al. (2013). Duodenal infusion of donor feces for recurrent Clostridium difficile. The New England journal of medicine 368, 407–415.

Warren, C.A., van Opstal, E.J., Riggins, M.S., Li, Y., Moore, J.H., Kolling, G.L., Guerrant, R.L., and Hoffman, P.S. (2013). Vancomycin treatment’s association with delayed intestinal tissue injury, clostridial overgrowth, and recurrence of Clostridium difficile infection in mice. Antimicrob Agents Chemother 57, 689–696.

Wilcox, M. (2015). Bezlotoxumab prevents clostridium difficile (C diff) infection recurrence: results of the MODIFY I trial (American Society for Microbiology).

Zaura, E., Brandt, B.W., Teixeira de Mattos, M.J., Buijs, M.J., Caspers, M.P., Rashid, M.U., Weintraub, A., Nord, C.E., Savell, A., Hu, Y., et al. (2015). Same Exposure but Two Radically Different Responses to Antibiotics: Resilience of the Salivary Microbiome versus Long-Term Microbial Shifts in Feces. MBio 6, e01693–01615.

