## Supplemental Figures for "The clinical drug candidate ebselen attenuates inflammation and promotes microbiome recovery after antibiotic treatment for *Clostridium difficile* infection"

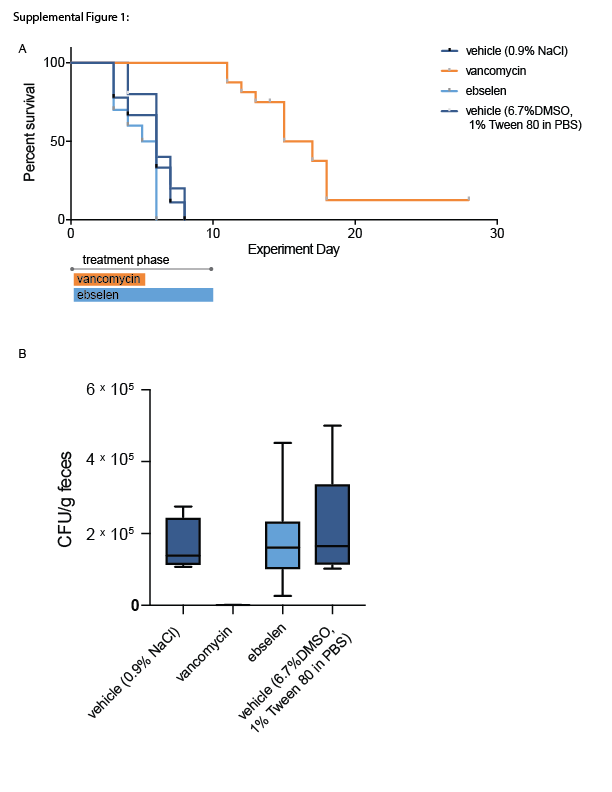

**Supplemental Figure 1: Hamster model of CDI**. A. Kaplan-Meier curve of hamster model of CDI (see Figure 1A for schematic). B. Quantification of *C. difficile* CFU/g feces collected from hamsters in (A) at day 3 post inoculation.

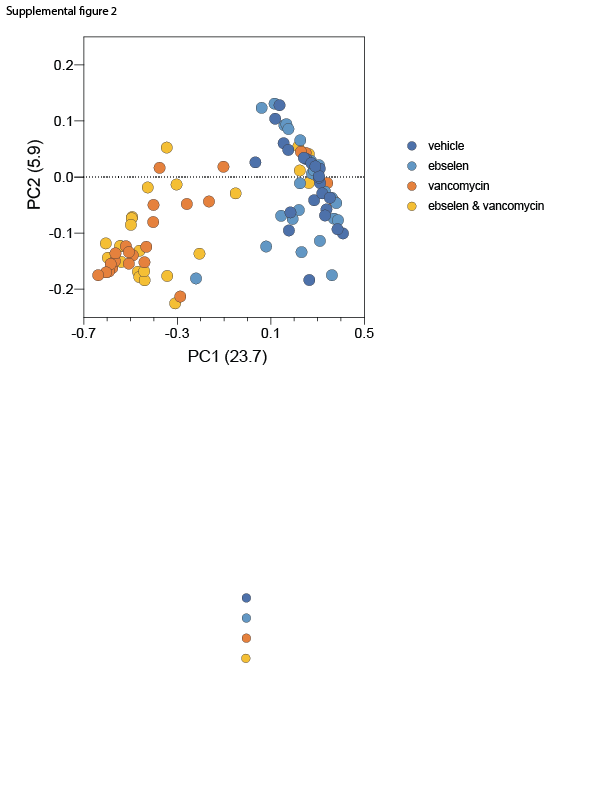

**Supplemental figure 2**: **5D healthy microbiome experiment**. PCoA plot of weighted UniFrac distances for healthy microbiome experiment (see Figure 2A for schematic) for GI microbiota days 0-5.

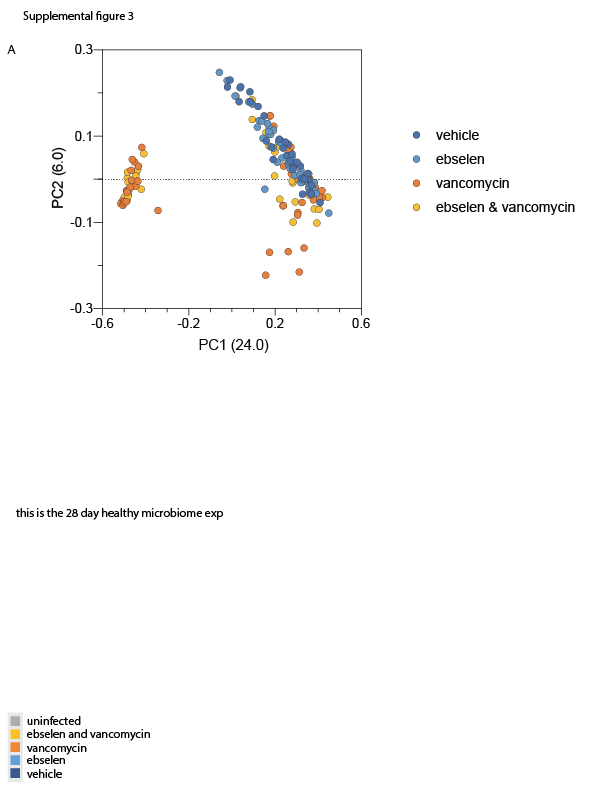

**Supplemental Figure 2: 28D healthy microbiome experiment.** PCoA plot of weighted UniFrac distances for healthy microbiome experiment (see Figure 3A for schematic) for GI microbiota over duration of experiment.

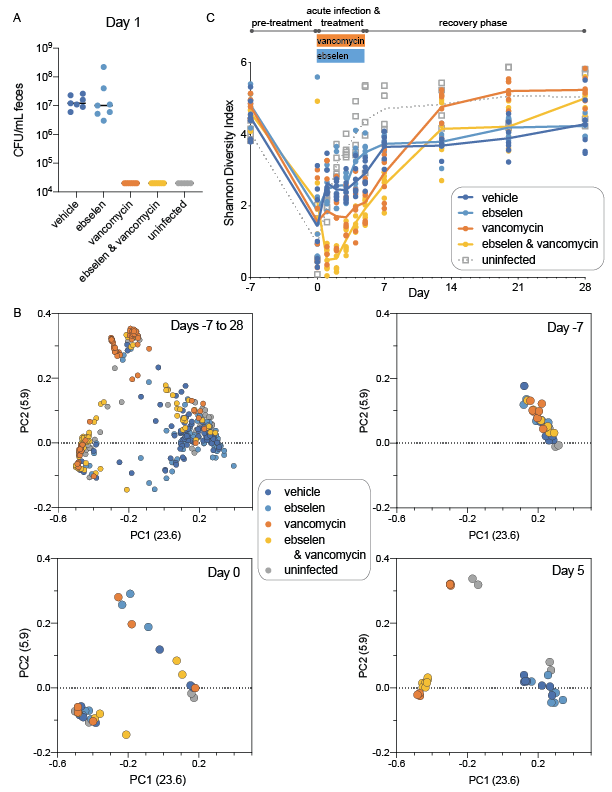

**Supplemental Figure 4: Mouse model of CDI**. A. Quantification of *C. difficile* CFU/mL feces collected from mice in mouse model of CDI (see Figure 4A for schematic) on day 1 post-inoculation. B. PCoA plot of weighted UniFrac distances for mouse model of CDI. Plots displayed for GI microbiota across all measured days of the experiment (top left) or days -7, 0, and 5 alone. C. Alpha diversity of GI microbial amplicon sequence variants (ASVs) measured over the pre-treatment, acute infection & treatment and recovery phases via Shannon diversity index.
